# Resveratrol induces H2A.X phosphorylation and H4K16 de-acetylation in *Toxoplasma gondii*

**DOI:** 10.1101/2020.08.25.266031

**Authors:** Susana M. Contreras, Agustina Ganuza, María M. Corvi, Sergio O. Angel

## Abstract

Resveratrol (RSV) is a multi-target drug that demonstrated activity against *Toxoplasma gondii* in macrophage and HFF cell line infection models. In addition to modulate redox homeostasis, RSV is also an activator of Sir2, a type III HDAC. RSV inhibited intracellular *T. gondii* tachyzoite growth at concentrations below the toxic effect on host cells. The IC_50_ value in a 24-hours treatment was 53 μM. After 96 hours of treatment the maximum non-toxic concentration for host cell, 20 μM, only inhibited *T. gondii* growth a 50%. RSV induced a reduction in H4K16 acetylation (H4K16ac), a mark associated to transcription, DNA replication and homologous recombination repair, without any effect on H3 acetylation (H3ac). RSV also enhanced the SQE motif phosphorylation on *T. gondii* H2A.X (termed γH2A.X), a DNA damage associated PTM. Sirtinol, a specific Sir2 inhibitor also inhibited *T. gondii* but did not altered the acetylation status of H3 and H4K16 as well as H2A.X phosphorylation. Our findings suggest a possible link between RSV and DNA damage or DNA repair process maybe due to DNA replication stress and/or another undetermined mechanism.

## Introduction

*Toxoplasma gondii* is an important pathogen for animals and human health, particularly during pregnancy and in immunocompromised patients [1]. The success of human infection is based on the ability of the tachyzoite to rapidly infect any nucleated cell. This event starts the asexual cycle which also occurs in other mammals and birds. The asexual phase is characterized by two stages, the rapidly replicating and highly disseminating tachyzoite, and the bradyzoite, which replicates slowly and is located in tissue cysts for the rest of the life of the animal or individual [2]. The replicative process can trigger a collapse in the DNA replication fork and its consequent double strand break (DSB). This effect could also be occurring at least in the *T. gondii* tachyzoite. On one hand, histone H2A.X is phosphorylated at the C-terminal SQEF motif to generate γH2AX foci in DSB sites. In *T. gondii*, it is possible to detect the presence of γH2AX under normal growth conditions [3, 4]. On the other hand, the ATM kinase inhibitor, key to triggers the DNA damage response (DDR), blocks H2A.X phosphorylation and stops the replication of *Toxoplasma* [4]. *Toxoplasma* possesses the conserved DSB repair machinery although it lacks some key effector proteins to decide the repair route: homologous recombination (HRR) or non-homologous end joining (NHEJ) [5]. Also, it has been observed that most of the components of the HRR pathway are essential and that a large number of drugs that are genotoxic or that inhibit DDR affect the growth of *Toxoplasma in vitro* and *in vivo* [6].

In addition to the phosphorylation of H2A.X and other proteins involved in DDR, there are other post-translational modifications of importance, including the ubiquitination of histones and acetylation of histone or non-histone proteins [5, 6]. Within acetylations, ATM kinase must be acetylated for its activation and histone H4 acetylated on its lysine in position 16 (H4K16ac) is a mark that facilitates the HRR pathway [6, 7].

Resveratrol (RSV; 3,5,4’-trihydroxystilbene) is a natural polyphenolic phytoalexin produced in plants. RSV is a multi-target drug, that among other effects can alter the activity of some histone deacetylases HDACI/II, DNA methyl-transferases (DNMT) and an activator of Sir2, a sirtuin belonging to HDACIII [6]. It has also been shown to be linked to the expression of genes associated with HRR, and enzymes which inhibits the HRR pathway [6]. Sir2 modulates the acetylation status of H4K16 [8, 9]. Interesting, RSV demonstrated opposite effects according to the dose, being that at low doses it is anti-oxidant and protects against DNA damage, while at higher doses it is pro-oxidant and generates DNA damage [10, 11].

In *in vivo* assays, it was shown that oral administration of RSV during acute *T. gondii* infection in mice would confer protection and survival [12] as well as moderate tissue inflammation and reduction of parasite replication [13]. Chen *et al*., [14] observed that incubation of extracellular tachyzoites with RSV during 24 hours affected their viability, probably by disturbing the redox homeostasis of the parasites. Of note, in addition to fork collapse, DNA can be damaged by oxidation. In addition, they observed that RSV reduced tachyzoite intracellular growth and promoted autophagy in infected macrophages. In parallel, Adeyemi *et al*., [15] identified RSV as a putative drug repurposing candidate against toxoplasmosis in a 62-compound screening. However, none of them analyzed the intracellular effect of RSV on *T. gondii-cell* tachyzoites.

In this work, we evaluated the effect of RSV on *T. gondii* growth and histone PTMs alteration. *T. gondii* expresses two type-III Sir2 histone deacetylases: TGME49_227020 and TGME49_267360, orthologues of *P. falciparum* Sir2A and Sir2B respectively and homologues to yeast Sir2 HDAC [16, 17]. The effect of RSV on H4K16 acetylation level was analyzed. In addition, changes in γH2A.X levels were also analyzed in control or treated intracellular tachyzoites. Collectively, our results show that RSV inhibited *T. gondii* growth and induced H4K16 de-acetylation. Finally, γH2A.X marks were highly enhanced in tachyzoites treated with RSV suggesting and association between this treatment and DSB damage.

## Materials and methods

### Parasite sources, culture and reagents

Tachyzoites of RH wild type and RH RFP (Red Fluorescent Protein) strains were cultured under standard conditions *in vitro* in immortal human foreskin fibroblasts (hTERT, ATCC^®^ CRL-4001) monolayers. RH RFP was kindly given by Dr. Silvia N. Moreno (University of Georgia, Athens, GA, USA). Cell monolayers were infected with tachyzoites and incubated with Dulbecco’s modified Eagle medium (DMEM, Invitrogen) high glucose supplemented with 10% (10X) or 1% (1X) of fetal bovine serum (FBS, Internegocios S.A., Argentina) and penicillin (10,000 units/ml) - streptomycin (10 mg/ml) solution (Gibco) at 37°C and 5% CO_2_ atmosphere. Resveratrol (Abcam,120726) solution was stocked at 10 mg/ml and sirtinol (Sigma, S7942) solution at 10 mM concentration. Both stocks were prepared in DMSO.

Commercial rabbit antibodies: anti-H3 (Abcam ab10799), anti-H3ac (Millipore, 06-599B), anti-H3K9me3 (Abcam, ab8898), anti-H4 (Abcam ab31830) and anti-H4K16ac (Abcam, ab109463). Alexa fluor secondary antibodies: goat anti-mouse 488 (Invitrogen, A11001) and anti-rabbit 595 (Invitrogen, A11037). Alkaline phosphatase–conjugated anti-rabbit or anti-mouse secondary antibodies (Santa Cruz Biotechnology).

Rabbit anti-*T. gondii* γH2A.X serum sample was obtained from Eurogentec (Belgium) on the basis of the peptide NH_2_-C+ GKHGV-S_(PO3H2)_-QEF −COOH, designed from *T. gondii* H2A.X amino acid sequence. Rabbit anti-rH2A.X was obtained from the Animal Facility at Facultad de Ciencias Exactas y Naturales (University of Buenos Aires, Argentina) on the basis of the recombinant rH2A.X [18]. Murine anti-SAG1 was kindly provided by Dr. Marina Clemente [19].

### Toxicity assay

To evaluate the cytotoxic effect of the drugs on host cells, fibroblasts (hTERT) were seeded at a 40% confluence (1.6 10^4^ cells/well) in 96-well plates for 24 hours. After this period, confluence was checked by microscopy and the medium replaced with fresh media containing different drugs at different concentrations. For experiments where cells were in contact with the drugs for 24 hours, the concentrations ranged between 25 to 200 μM. For experiments in which the cells were in contact with the drugs for 96 hours, the concentrations tested ranged between 97 nM to 200 μM. Then, in both cases, cytotoxicity was determined by reduction of 3 (4,5 dimethyl-2-thiazoyl)-2,5-diphenyltetrazole bromide (MTT). The absorbance was measured from the bottom of the plate at 540 nm using a BioTek Synergy plate reader. Graphs indicate the relative viability of the cells with respect to the control with DMSO 0.5% in culture media (100% viability).

### Growth assay

Freshly lysed RH RFP parasites were used to measure tachyzoite replicative growth capacity by direct fluorescence in presence or absence of drugs [20]. For intracellular exposition, hTERT cells were seeded at 40% confluence (1.6 10^4^ cells/well) on 96-well plates for 24 hours. After this period, confluence was checked by microscopy and half of the plate was infected with 10,000 RH RFP parasites. The other half of the plate was used as control (host-cells + different concentrations of drugs tested). After 3 hours of infection, the medium of the whole plate was replaced by fresh media 1X containing increasing concentrations of drugs or vehicle. Parasites were then allowed to replicate for 24 or 96 hours on host cells. Before fluorescence analysis, infected monolayers were observed by inverted microscope to check hTERT cell integrity. The RFU (<u>R</u>elative <u>F</u>luorescent <u>U</u>nit) of the RFP data was collected using a Synergy H1 Hybrid Multi-Mode Microplate Reader (Biotek) in fluorescence mode, exciting at 540 nm and measuring the red fluorescence emission at 590 nm. Each concentration of drug was measured in triplicates, in three different experiments with similar results. The basal fluorescence was estimated in 234 +/- 2.3 RFU. We also calculated the IC_50_ value for each drug. A nonlinear fit of log of drug concentration versus normalized response (control equal 100%) was carried out. The data was analyzed using Microsoft Excel and then extrapolated to the GradPad Prism 6 program (version 6.1). IC_50_ was obtained by GraphPad Prism 6: data were normalized with 0 as the smallest value and transformed to semi-logarithmic scale (x=log(x)). After that, they were analyzed as a nonlinear regression parameter-dose-response inhibition-log(inhibitor) vs normalized response-variable slope.

### Indirect immunofluorescence assay (IFA)

Intracellularly treated tachyzoites were incubated during 24 hours with drugs, washed with PBS 1X and fixed with methanol for 10 min at −20°C. Then the cells were permeabilized with 0.2% v/v Triton X-100 in PBS for 10 min and blocked with 3% w/v BSA in PBS for 30 min at room temperature (r.t.). After this, appropriate primary antibodies were added diluted in 0.5% w/v BSA in PBS for 60 min at r.t. Following incubation, cells were washed three times with PBS and then incubated with the corresponding alexa fluor-conjugated secondary antibodies for 60 min at r.t. Samples were mounted on coverslips with mounting medium (Mowbiol) containing 10 μg/μl of DAPI (4,6-diamidino 2-phenyl-indole) to stain the nuclei. Primary antibodies were diluted 1:200 whereas secondary antibodies were diluted 1:2,000. Samples were analyzed by epifluorescence microscopy (Zeiss Axioscope) equipped with a 63X objective and Zeiss Axiocam 506 mono microscope camera. Images and fluorescence intensity analysis were obtained with Fiji (ImageJ) program [21].

### Western blot

Proteins from purified parasites (10^7^ parasites/lane) from each treatment were resolved by 15% SDS-PAGE and transferred onto a PVDF (Polyvinylidene Difluoride) membrane. Non-specific binding sites were blocked with 5% non-fat-dried milk in TBS containing 0.1% v/v Tween-20 (TBS-T) for 6 h and the membranes were then incubated overnight with primary antibodies. The following primary antibodies and corresponding dilutions were: mouse anti-H3 (1:250), rabbit anti-H3ac (1:400), mouse anti-H4 (1:250), rabbit anti-H4K16ac (1:500), rabbit anti-γH2AX (1:100), rabbit anti-H2AX (1:5.000), mouse anti-SAG1 (1:500). The membranes were then washed several times with TBS-T prior to incubation with alkaline phosphatase-conjugated anti-rabbit or anti-mouse secondary antibodies, diluted 1:10,000. Immunoreactive protein bands were visualized by the NBT-BCIP method (Sigma-Aldrich™ Argentina S.A). Intensities of bands were quantified from scanned images using ImageJ software and the value obtained from each band was normalized by the loading control SAG1 and then each histone PTM was normalized by their respective histone (H3 or H4).

### Statistics

Data were expressed as mean ± SD from three to four different experiments. The variations of the data were analysed with the GraphPad Prism (6.1) software, using ordinary one-way ANOVA and Tukey’s multiple comparison test (* p ≤ 0.05, **p ≤ 0.01, *** p ≤ 0.001 and ****p ≤ 0.0001) (GraphPad Prism 6.1). Unpaired Student’s *T* test was also used.

## Results and Discussion

### Toxicity assay on hTERT monolayer and on tachyzoite growth

RSV has already demonstrated its anti-*T. gondii* effect *in vitro* in two recent works [14, 15]. In order to confirm these results in our study model, uninfected or infected hTERT host cell was incubated with different doses of RSV and in two incubation periods (24 and 96 hours). In the 24-hours incubation experiments, 80% of the cells remain viable at concentrations of 100 μM, while after 96 hours of exposure, 90% remain viable at 12.5 μM (Fig 1). In order to test the impact of RSV on *T. gondii* lytic cycle, transgenic RH RFP (<u>R</u>ed <u>F</u>luorescent <u>P</u>rotein) tachyzoites were intracellularly grown during 24 and 96 hours in presence of RSV. A direct relationship between the amount of tachyzoites and the amount of fluorescence emitted by those parasites was assumed. As it can be observed, RSV showed a dose-dependent effect at 24 hours in intracellular *T. gondii* growth (Fig. 1) with an IC_50_ value of 53 +/- 4 μM. This result agrees with those observed by Chen et al., [14], in which a 50 μM of RSV blocked *T. gondii* intracellular growth. When intracellular tachyzoites were grown for 3 hours under normal conditions and were then treated for 96 hours with different doses of RSV, the maximum dose of 20 μM only reduced a 50% the tachyzoite growth rate (Fig. 1). Adeyemi *et al*., [15] identified RSV as one of the 62 compounds with anti-*T. gondii* effect with an IC_50_ value of 1.03 μg/ml (4.4 μM) after 72 hours of incubation. The authors also observed that HFF cells were 100% viable at 2 μg/ml (8.5 μM). Taken bibliographic and our results together, RSV is a natural product with potential for an alternative *Toxoplasma* infection therapy maybe by combination with other drugs.

**Fig 1.**
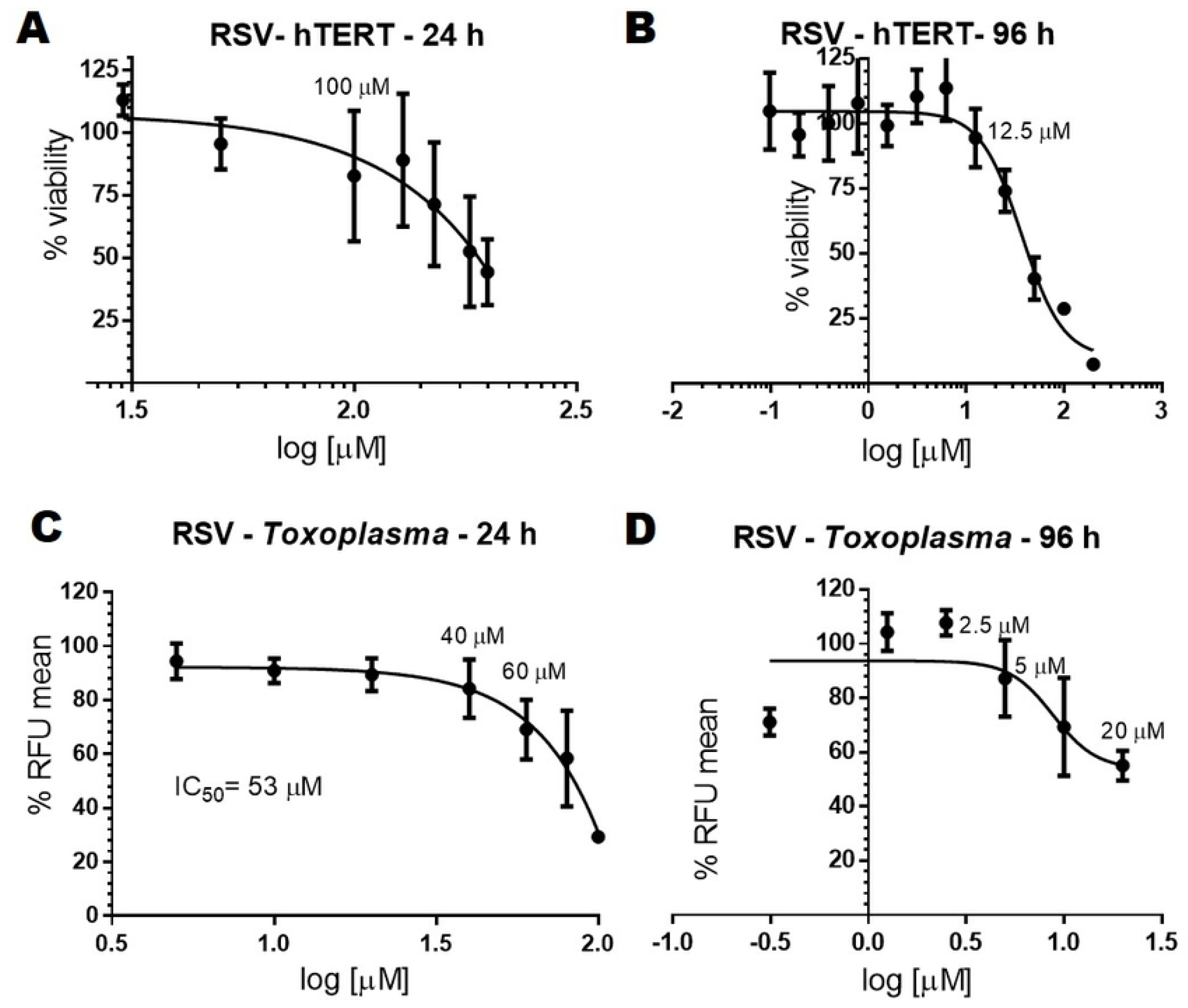
Effect of resveratrol (RSV) on cell viability and *T. gondii* growth. **A,B**. hTERT cell monolayers were incubated for 24 (**A**) or 96 hours (**B**) in the absence or presence of RSV. Cytotoxicity was estimated by 3-(4,5-dimethylthiazol-2-yl)-2,5-diphenyl-tetrazolium bromide reduction (Absorbance at 540 nm) which is presented as relative to the untreated controls (defined as 100% viability). Determinations were performed on triplicate. The results are representative of three independent experiments. Untreated controls contained 0.5% v/v DMSO. **C, D.** Effect of resveratrol on intracellular RH tachyzoite growth. A dose-dependent growth curve relative to RFP fluorescence after 24 hours (**C**) or 96 hours (**D**) treatment is shown. IC_50_ value was determined only for 24 hours because at 96 hours the dose could not be increased above 20 μM. Three independent experiments performed in triplicate each time and the data are presented as mean ± standard deviation (SD).

### Effect of RSV and sirtinol on H3 and H4 post-translational modifications in intracellular tachyzoites

RSV is a possible activator of *T. gondii* Sir2A and/or Sir2B, among other targets, an enzyme associated with regulating the de-acetylation status of histones H3 and H4. H3ac marks were observed in nuclei of intracellular tachyzoites with similar intensities in treated or untreated tachyzoites (DMSO, control condition) (Fig. 2A). In fact, Western blot analysis did not show significant differences in band intensities (Fig. 2B). Of note, the antibody used is against H3K9acK14ac marks. Maybe, only one of these marks are being modulated by some of *T. gondii* Sir2. Unfortunately, the anti-H3K9me3 used here did not work in immunofluorescence assay (Fig. 2C). Sirtinol (Sir two inhibitor naphthol) is a sirtuin inhibitor, highly specific for human SIRT1 and SIRT2 with IC_50_ of 131 μM and 38 μM respectively [22]. We decided to include this drug to see the opposite effect of resveratrol. Based on a viability study on hTERT cell (S1 Fig.), the concentration used infected cells was 100 μM. Sirtinol did not present any effect on H3ac status (Fig 2A). However, sirtinol affected *T. gondii* growth in a dose dependent manner with an IC_50_: 27 +/- 2.5 μM (S1 Fig.).

**Fig 2.**
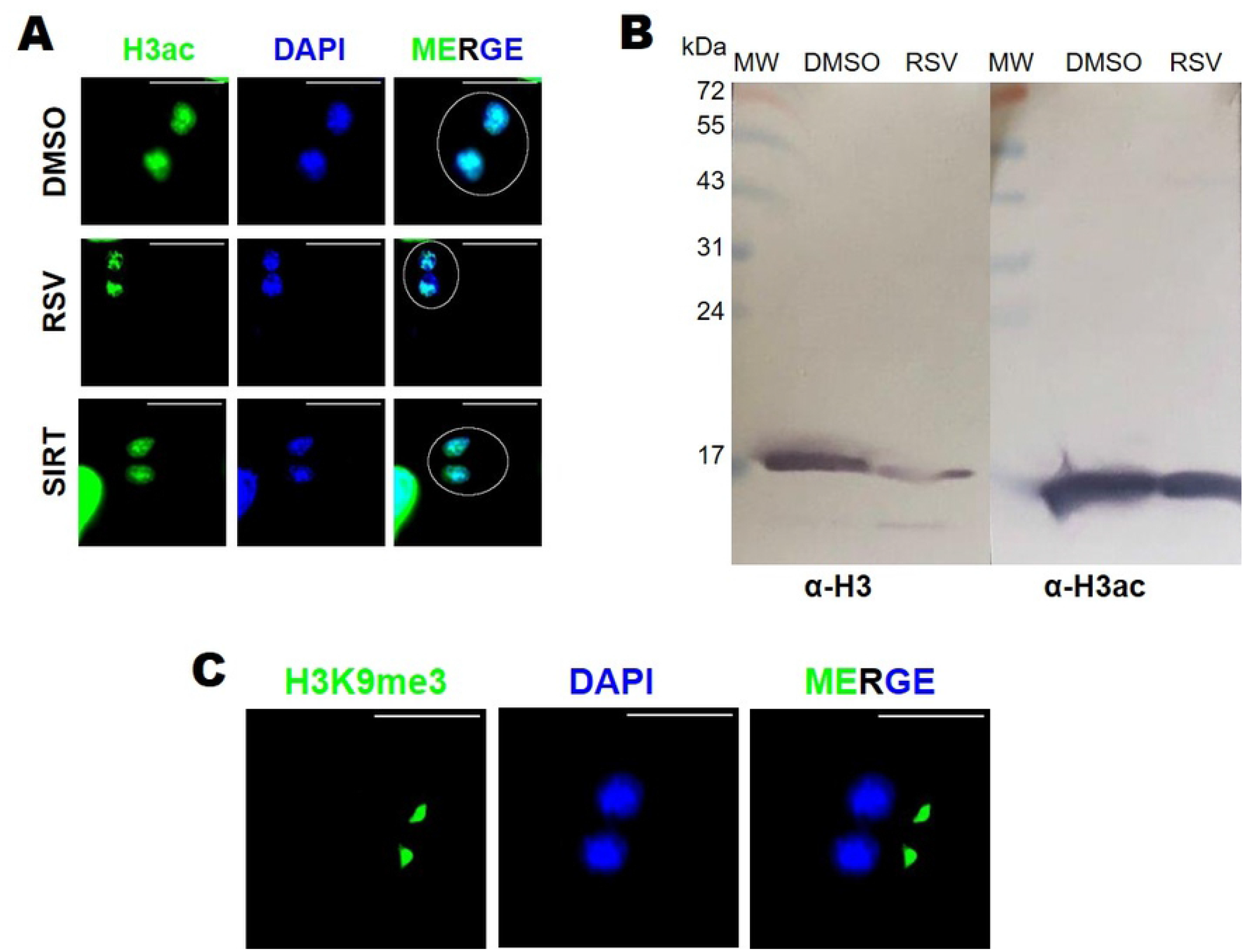
Effect of intracellular exposition to Resveratrol (RSV) on H3 post-translational acetylation. **A.** Indirect immunofluorescence with anti–H3Ac antibody (1:200). Two parasites per PV (white circle) is shown. The upper right white lines correspond to the scale bar: 10 μm. Nuclei were stained with DAPI. Intracellular tachyzoites were treated with RSV (100 μM), Sirtinol (100 μM) or DMSO 0.5% during 24 hours. **B**. Western blot of purified *T. gondii* tachyzoite lysates with anti-H3 (1:250) or anti-H3ac (1:400). Lysates were obtained from purified intracellular tachyzoites previously treated with RSV (50 μM), Sirtinol (50 μM) or DMSO 0.5% during 24 hours. **C.** Testing of anti-H3K9me3 (1:200) antibody under normal conditions by indirect immunofluorescence. Panels are representative of three independent experiments with similar results

Tachyzoites treated with RSV presented a significant weaker H4K16ac labelling in comparison with DMSO or sirtinol treatment (Fig. 3A and S2 Fig). This result was confirmed by Western blot analysis, in which RSV reduced the H4K16ac mark whereas sirtinol did not show significant differences when compared to DMSO control (Fig. 3B). The weak signal observed by H4K16ac in presence of RSV was evident. This result is in agreement with another recent study in which RSV treatment decreased H4K16ac levels [23]. In addition, in yeast, Sir2 negatively controlled the activation of DNA replication origins within heterochromatin and euchromatin by de-acetylating H4K16 [24]. Whether RSV is affecting *T. gondii* replication via H4K16 de-acetylation requires of further studies.

**Fig 3.**
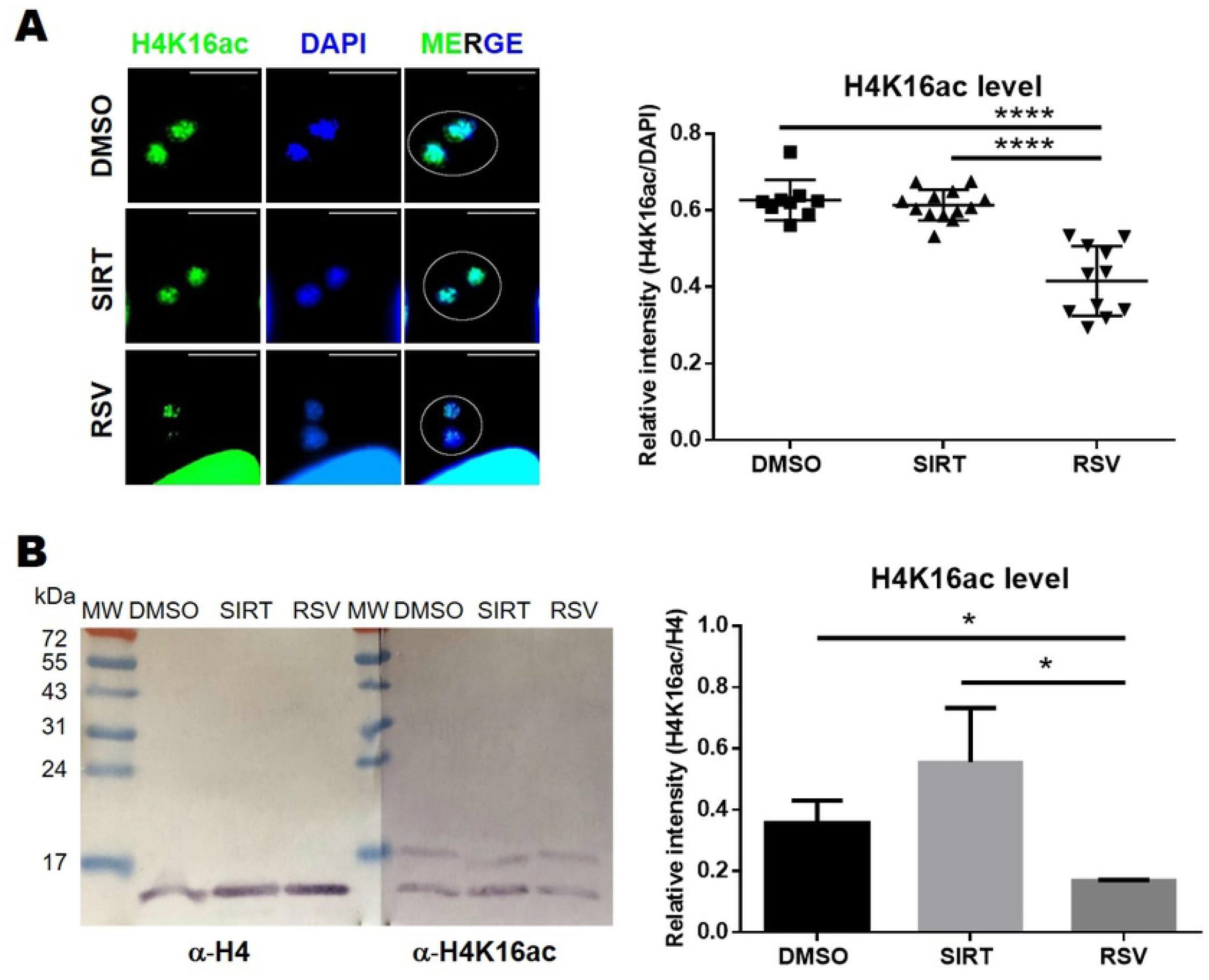
Effect of intracellular exposition to resveratrol (RSV) on H4 post-translational acetylation. **A.** Indirect immunofluorescence with anti-H4K16ac antibody (1:200). Two parasites per PV (white circle) is shown. The upper right white lines correspond to the scale bar: 10 μm. Nuclei were stained with DAPI. Intracellular tachyzoites were treated with RSV (100 μM), Sirtinol (100 μM) or DMSO 0.5% during 24 hours. DAPI and fluorescence intensities were quantified from 10 nuclei (S2 Fig) and plotted as relative intensity. The panel and graph are representative of three independent experiments with similar results. **B**. Western blot of *T. gondii* lysates with anti-H4 (1:250) or anti-H4K16ac (1:500). Lysates were obtained from purified intracellular tachyzoites treated with RSV (50 μM), Sirtinol (50 μM) or DMSO 0.5% during 24 hours. H4K16ac band intensities were quantified and relativized to H4 histone and plotted. Statistical analysis was performed with one-way ANOVA and Tukey’s multiple comparison test (* p ≤ 0.05 and ****p ≤ 0.0001).

### Effect of RSV and sirtinol on H2A.X phosphorylation in intracellular tachyzoites

The presence of H4K16ac in DNA damaged by DSB is able to recruit DNA repair proteins associated to the HRR pathway [7], among them ATM kinase that initiate the DNA damage response [6]. ATM-like kinase activity was detected in *T. gondii* [4]. In order to determine if the effect of RSV and sirtinol is associated to DSB damage, the γH2A.X level (DSB level mark) was tested by Western blot. A specific rabbit anti-*T. gondii* phosphorylated peptide in the SQE motif of *T. gondii* H2A.X C-terminal end was prepared (see materials and methods). To confirm its specificity, a Western blot against the recombinant non-phosphorylated *T. gondii* H2A.X was performed. As it can be observed, the rabbit anti-TgγH2A.X antibody does not recognize *T. gondii* H2A.X (Figure 4A), but reacts with an expected band of 14.5-kDa in *T. gondii* lysate (Fig 4B). After treatment with RSV, the γH2A.X signal is increased compared to DMSO control, whereas sirtinol treatment produced a mild effect, below a 2-fold enrichment (Fig. 4B).

**Fig 4.**
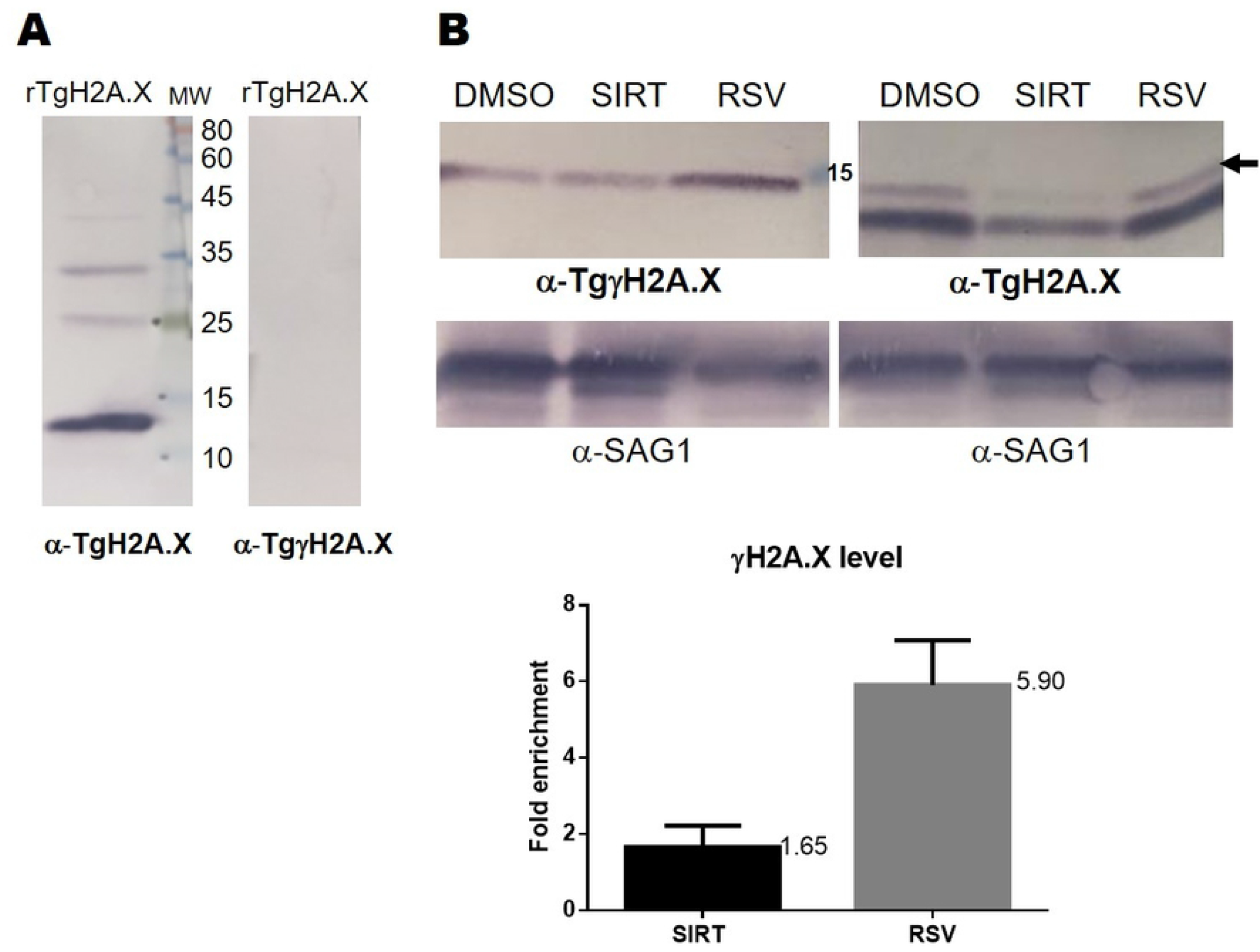
Effect of intracellular exposition to resveratrol (RSV) on *T. gondii* H2A.X phosphorylation (γH2AX). **A**. Western blot of *T. gondii* recombinant H2A.X (rH2A.X; 200 ng/lane) expressed in *Escherichia coli* and purified by nickel resin. Rabbit anti-rTgH2A.X (α-TgH2A.X, 1:5,000) and anti-*T. gondii* phosphorylated peptide (α-TgγH2A.X, 1:100). Phosphorylated peptide sequence was NH_2_-C+ GKHGV-S_(PO3H2)_-QEF −COOH. **B.** Western blot of *T. gondii* lysates with anti-SAG1 (*T. gondii* surface antigen 1, 1:500), anti-TgH2A.X or -anti-TgγH2A.X. Lysates were obtained from purified intracellular tachyzoites previously treated with RSV (50 μM), Sirtinol (50 μM) or DMSO 0.5% during 24 hours. H2A.X (arrow) and γH2A.X band intensities were quantified and normalized against SAG1 band intensities. After that, relative intensity bands (γH2A.X/H2A.X) were calculated for each treatment. Normalized signal to SAG1 were calculated from relative values in comparison to DMSO control. Results are mean of three replicates ± SD.

Here we observed that the γH2A.X level is increased in presence of RSV 50 μM, a mark compatible with DNA damage. In another model, RSV at 50 μM induced DNA damage, S-phase arrest and enhanced γH2AX levels in a panel of head and neck squamous cell carcinoma lines [25]. In addition, RSV was associated to HRR inhibition through the inactivation of tyrosyl-tRNA synthetase, an enzyme associated to protein synthesis but also to HRR gene regulation when translocate to nucleus [26, 27]. If that were the case, RSV could be used in combination with a drug that inhibits the DSB repair pathway in *T. gondii*, as previously suggested [6].

These analyzes suggest that RSV would impact on *T. gondii* histone H4K16ac, the histone mark associated to transcription activation, DNA replication and DSB repair, and increase phosphorylation of H2A.X. By contrast, the anti-*T. gondii* mechanism of sirtinol could not be elucidated. Interestingly, *P. falciparum* Sir2A, an orthologue of *T. gondii* Sir2A did not show to be sensitive to sirtinol *in vitro* but it was inhibited by nicotinamide [28], a drug that had not effect on *T. gondii* growth [29]. An effect on host cell metabolism due to sirtinol that perturb *T. gondii* growth cannot be ruled out. In fact, in a model of cancer *in vitro* it was observed that sirtinol presented an anti-proliferative and apoptotic effect via the transcription factor FOXO3a and the kinase AKT [30]. Inhibition of host cell PI3K/AKT signaling pathway reduced *T. gondii* proliferation [31].

## Conclusions

In summary, we have observed that RSV and sirtinol are effective drugs against *T. gondii* growth *in vitro*. Whereas the biological process in which sirtinol produces its effect remains intriguing, RSV showed to alter parasite growth and reduce H4K16ac level, a PTM associated with gene transcription, heterochromatin spread, DNA replication and/or DNA damage repair. Interestingly, RSV also enhances the phosphorylation of SQE motif on *T. gondii* H2A.X, suggesting a link between the drugs with DNA damage and/or DNA replication stress. Our results add new evidence of putative effect of the multi-target drug RSV in addition to the redox homeostasis alteration suggested recently [14]. Finally, we considered that RSV could be an interesting drug to analyze chromatin modulation in *T. gondii* as well as novel candidate for alternative *T. gondii* infection treatment.

## Author Contributions

Conceptualization: Sergio O. Angel

Data curation: Susana M. Contreras, Agustina Ganuza, Maria M. Corvi

Formal analysis: Susana M. Contreras, Maria M. Corvi, Sergio O. Angel

Investigation: Susana M. Contreras, Sergio O. Angel

Methodology: Susana M. Contreras, Agustina Ganuza, Maria M. Corvi, Sergio O. Angel

Resources: Sergio O. Angel

Supervision: Maria M. Corvi, Sergio O. Angel

Validation: Susana M. Contreras, Maria M. Corvi, Sergio O. Angel

Visualization: Susana M. Contreras, Maria M. Corvi, Sergio O. Angel

Writing – original draft: Sergio O. Angel.

Writing – review & editing: Susana M. Contreras, Maria M. Corvi, Sergio O. Angel

## Data Availability Statement

All relevant data are within the manuscript and its Supporting Information files.

## Competing interests

The authors have declared that no competing interests exist.

## Funding

This work was supported by the Ministerio Nacional de Ciencia y Tecnología (MINCyT, PICT 2015 1288) and Consejo Nacional de Investigaciones Científicas y Tecnológicas (CONICET, PIP 11220150100145CO) and by National Institute of Health (NIH-NIAID 1R01AI129807).

## Acknowledgments

SM Contreras (Fellow), MM Corvi (Researcher) and SO Angel (Researcher) are member of CONICET. SO Angel (Full) and MM Corvi (Adjunct) are Professors of Universidad Nacional General de San Martin (UNSAM). We thank Agustina Ganuza (Comisión de Investigaciones Científicas, CIC) for her valuable help in the interpretation and analysis of FACS data and Laura Vanagas for critically reading the manuscript.

## Conflicts of Interest

The authors declare no conflict of interest.

## Ethics approval and consent to participate

Not applicable.

---

**S1 Fig. Effect of sirtinol on cell viability and *T. gondii* growth. A.** hTERT cell monolayers were incubated for 24 hours in the absence or presence of sirtinol. Cytotoxicity was estimated by 3-(4,5-dimethylthiazol-2-yl)-2,5-diphenyl-tetrazolium bromide reduction (Absorbance at 540 nm) which is presented as relative to the untreated controls (defined as 100% viability). Determinations were performed on triplicate. The results are representative of three independent experiments. Untreated controls contained 0.5% v/v DMSO. **B.** A dose-dependent growth curve relative to RFP fluorescence after 24-hour treatment is shown. Three independent experiments performed in triplicate each time and the data are presented as mean ± standard deviation (SD). IC_50_ value was determined by GraphPad Prism 6.1.

**S2 Fig. Quantification of H4K16ac mark with or without drug treatment.** Panel shows 10 nuclei of intracellular tachyzoites under RSV, SIRT treatment or DMSO control, labelled by Indirect immunofluorescence with α-H4K16ac (1:200). Nuclei were also stained with DAPI. Antibody fluorescence and DAPI signals were quantified and plotted in different graph (see Fig 3).

